# Characterisation and modelling of a polyhydroxyalkanoate synthase from *Aquitalea* sp. USM4 reveals a mechanism for polymer elongation

**DOI:** 10.1101/235556

**Authors:** Aik-Hong Teh, Nyet-Cheng Chiam, Go Furusawa, Kumar Sudesh

**Affiliations:** Centre for Chemical Biology, Universiti Sains Malaysia, 10 Persiaran Bukit Jambul, Bayan Lepas, 11900 Penang, Malaysia; Ecobiomaterial Research Laboratory, School of Biological Sciences, Universiti Sains Malaysia, 11800 Penang, Malaysia

**Keywords:** polyhydroxyalkanoate, PHA synthase, enzyme mechanism, substrate specificity, homology modelling, ligand docking

## Abstract

Polyhydroxyalkanoate synthase, PhaC, is a key enzyme in the biosynthesis of PHA, a type of bioplastics with huge potential to replace conventional petroleum-based plastics. While two PhaC structures have been determined recently, the exact mechanism remains unclear partly due to the absence of a tunnel for product passage. The PhaC from *Aquitalea* sp. USM4, PhaC*_Aq_*, was characterised and showed a *K*_m_ of 394 µM and a *k*_cat_ of 476.4 s^−1^ on the 3HB-CoA substrate. A model based on the structure of the closely related PhaC from *Cupriavidus necator*, PhaC*_Cn_* revealed a three-branched tunnel at the dimeric interface. Two of the branches open to the solvent and serve as the putative routes for substrate entrance and product exit, while the third branch is elongated in a PhaC1 model from *Pseudomonas aeruginosa*, indicating a function of accommodating the hydroxyalkanoate (HA) moiety of the HA-CoA substrate. Docking of the two tetrahedral intermediates formed during catalysis suggests a PHA elongation mechanism that requires the HA moiety of the ligand to rotate ~180°. Both classes I and II PhaCs share a common mechanism for polymer elongation, and substrate specificity is determined in part by a bulky Phe/Tyr/Trp residue in the third branch in class I, which is conserved as Ala in class II to create room for longer substrates. The PhaC*_Aq_* model provides fresh insights into a general PhaC mechanism, pinpointing key residues for potential engineering of PhaCs with desirable characteristics.

## 1. Introduction

Polyhydroxyalkanoates (PHAs) are polyesters synthesised by a wide range of bacteria and archaea as carbon storage, usually when carbon sources are in excess while other essential nutrients such as nitrogen and phosphorous are limited [1]. Due to their biodegradability and biocompatibility, PHAs have attracted substantial interest not only as a sustainable alternative to conventional petroleum-based plastics [2], but also for applications in specialty markets such as medical devices. PHAs are accumulated in intracellular granules, and many proteins are localised to the surface of these granules. One of them is the key enzyme for PHA biosynthesis, PHA synthase, which polymerises hydroxyalkanoates (HAs) from HA-CoA and releases CoA during the reaction [3,4].

Four classes of PHA synthase, or PhaC, have been reported based on the enzyme’s substrate specificity and subunit composition [3,5]. PhaCs from classes I and II form homodimers, whereas those from classes III and IV form heterodimers with PhaE and PhaR respectively. Despite a relatively high identity of ~40% between the first two classes, class I PhaCs usually are reported to incorporate monomers of short chain length (C3–C5), while class II PhaCs can also incorporate monomers of medium chain length (C6–C14). Nevertheless, the strict conservation of an Asp–His–Cys catalytic triad in all PhaCs and a lag phase in CoA release observed in many of them suggest a common catalytic mechanism shared by all four classes.

Recently the dimeric structures of class I PhaCs from *Cupriavidus necator* [6,7], PhaC*_Cn_*, and from *Chromobacterium* sp. USM2 [8], PhaC*_Cs_*, have been solved. The PhaC*_Cn_* structure is in a partially open form with an obvious channel proposed for substrate entrance, while the PhaC*_Cs_* structure is in a closed form with no visible access to the catalytic site. Both differ in their dimer arrangement with a large conformation change in the so-called LID region (Supplementary Fig. S1), which is highly flexible and involved in dimerisation. However, due to the absence of a distinct tunnel for product passage, the exact mechanisms for polymer elongation and substrate specificity remain unknown.

From the PhaC*_Cn_* structure, two different mechanisms that utilise a single active site for PHA biosynthesis have been proposed. The first suggests a substrate entrance which is also used for the release of free CoA, and a separate egress route for the elongation of the nascent PHA chain [6]. The second, on the other hand, advocates a ping-pong mechanism in which the substrate and the growing PHA chain take turn to enter and leave the catalytic site through a single tunnel, with the catalytic Cys acting to transfer one HA monomer from HA-CoA to the PHA chain every cycle [7]. While the first model is consistent with the detection of a product covalently bound to the catalytic Cys [9], the proposed egress route was too narrow and would require substantial conformational changes for the growing PHA chain to pass through [6].

In an attempt to gain further insights into the PhaC mechanism, we have characterised and constructed a model of a class I PhaC from *Aquitalea* sp. USM4 (JCM 19919), PhaC*_Aq_*, which can produce PHAs containing 3-hydroxyvalerate (3HV), 4-hydroxybutyrate (4HB) and 3-hydroxy-4-methylvalerate (3H4MV) [10]. From the PhaC*_Aq_* model, we have identified a tunnel that traverses the dimer interface, providing a possible explanation for a chain elongation mechanism, substrate specificity, as well as the occurrence of the lag phase. While further structural studies are needed to investigate the binding patterns of the substrate and product, the PhaC*_Aq_*’s model has yielded fresh insights into a probable universal mechanism for PHA biosynthesis.

## 2. Materials and methods

### 2.1. Cloning, protein expression and purification

The gene encoding PhaC*_Aq_* was amplified by PCR using primers containing the NdeI and BamHI restriction sites, with a His-tag sequence inserted before the stop codon. The PCR product was ligated into the pET-21a vector (Novagen), and transformed into *Escherichia coli* BL21(DE3). The cells were grown in LB medium supplemented with 100 µg/ml ampicillin at 37 °C, and expression was induced with a final concentration of 100 µM isopropyl-*β*-D-thiogalactopyranoside (IPTG) when the OD_600_ reached 0.6. After seven hours of incubation at 25 °C, the cells were harvested by centrifugation and resuspended in 50 mM Tris-HCl, 300 mM NaCl, 5% glycerol, pH 7.4, lysed by sonication and centrifuged to remove cell debris. The supernatant was loaded onto a Ni^2+^-charged Protino Ni-IDA column (Macherey-Nagel), and PhaC*_Aq_* was eluted stepwise with up to 1.0 M imidazole. The Ni^2+^ affinity chromatography was repeated twice, followed by size-exclusion chromatography on a Superdex 200 column (GE Healthcare). Protein concentration was determined using the Bradford assay.

### 2.2. Enzyme activity assays

Enzyme activity of PhaC*_Aq_* was determined in triplicate in a discontinuous approach by measuring the formation of 3-thio-6-nitrobenzoate (TNB) dianion at 412 nm, which corresponded to CoA release. The assay was carried out at 30 °C in a 100 μl mixture of 150 mM potassium phosphate buffer (pH 7.2), 0.2% (w/v) glycerol, 0.005 % (w/v) Triton X-100, 0.2 mg/mL BSA, 3 nM enzyme, and 0.2–4 mM 3HB-CoA. At various intervals, 20 μl reaction mixture was removed and quenched with 50 μl of chilled 10% trichloroacetic acid (TCA). After centrifugation, 65 μl supernatant was mixed with 385 μl of 500 μg/mL 5,5′-dithiobis(2-nitrobenzoic acid) (DTNB). The kinetic constants, *K*_m_ and *k*_cat_, were determined from the Lineweaver–Burk plot using a molar absorption coefficient of 13.6 mM^−1^ cm^−1^ for TNB.

### 2.3. Modelling of PhaCs and ligand docking

All PhaC models (PhaC*_Aq_*, PhaC*_Cs_*, PhaC1 *_Pa_* and PhaC2*_Pa_*) were constructed with the Rosetta modelling software [11], using the PhaC*_Cn_* structure (PDB ID: 5HZ2) as a template. Ligands were docked manually into the catalytic site followed by geometry minimisation with the Phenix software [12], allowing only the side chains to rotate while the main chains were fixed. The two tetrahedral intermediates were generated by docking the CoA moiety of a 3HDD-CoA molecule, the Cys319-bonded (3HV)_5_ chain, and the 3HDD moiety respectively into branches I, II, and III. Orientation and position of the ligands for the other steps were then determined based on the two tetrahedral intermediates. All structures were rendered with PyMol (https://pymol.org).

### 2.4. Phylogenetic tree of PhaCs

A total of 43 PhaC sequences were retrieved from the GenBank database (https://www.ncbi.nlm.nih.gov/genbank/) and aligned using MUSCLE [13]. The esterase sequence from *Sulfolobus solfataricus* strain P1 was added as an outgroup. A phylogenetic tree was constructed using neighbour-joining methods [14] implemented in MEGA7 [15] with 1000 bootstrap replications.

## 3. Results and discussion

### 3.1. Large oligomerisation of PhaC_Aq_

Full-length PhaC*_Aq_* of 595 residues existed predominantly as large oligomers, eluting close to the void volume during size-exclusion chromatography with an estimated huge size of about 1 MDa (Fig. 1). In contrast, PhaC*_Cn_* was reported to exist in an equilibrium between monomer and dimer [16]. As both synthases share a comparatively high identity of 47%, the large oligomerization of PhaC*_Aq_* could have been due to regions not well conserved between them, i.e. the N-terminal domain whose first 98 residues of both synthases are poorly conserved (Supplementary Fig. S1).

**Fig. 1.**
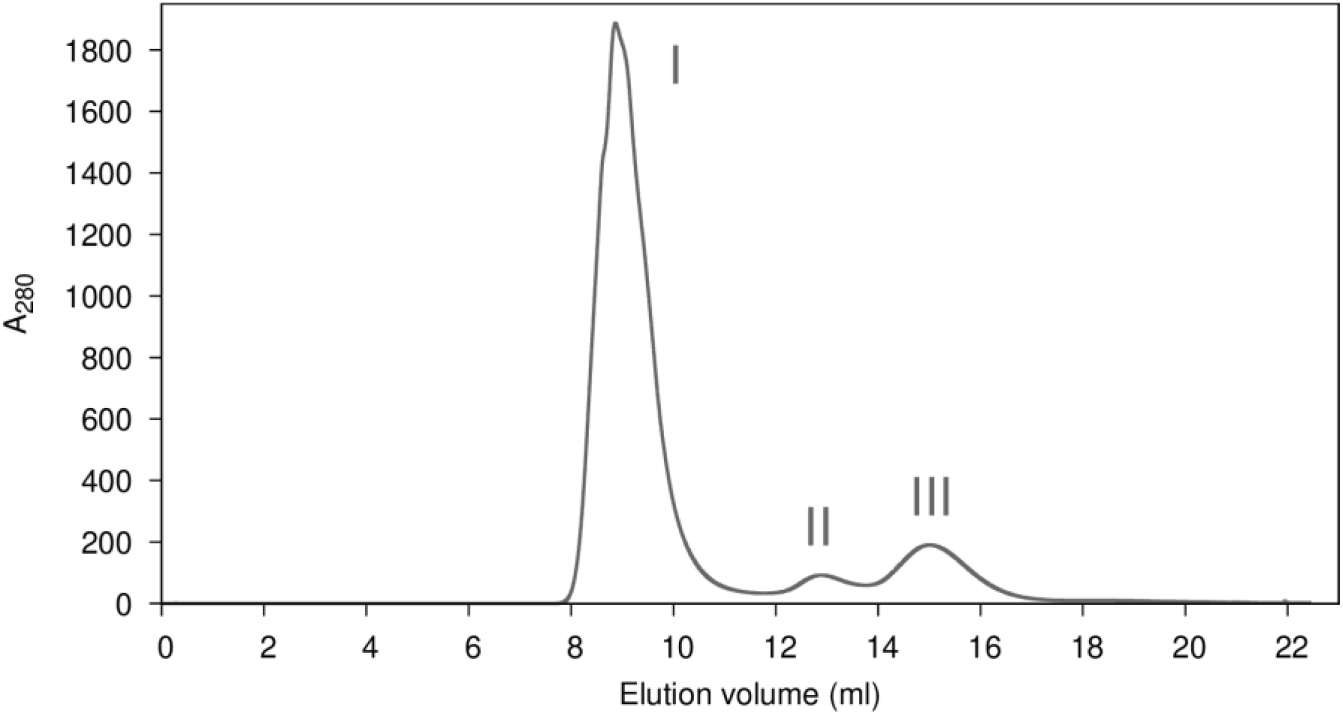
Size-exclusion chromatography. PhaC*_Aq_* eluted predominantly as large oligomers (I), and only two small fractions as dimers (II) and monomers (III).

The N-terminal domain reportedly plays an important role, as deleting PhaC*_Cn_*’s first 88 residues led to a significantly reduced activity [17]. PhaC*_Cn_*’s N-terminal domain was also found localized to PHA granules when co-expressed with full-length PhaC*_Cn_* and PhaAB in an *E. coli* strain [18], suggesting a role in anchoring the synthase to the granules. Interestingly, PhaC*_Cs_* was also found to exist in a monomer–dimer equilibrium despite a high identity of 78% with PhaC*_Aq_*, but it lacks the first 29 residues present in the latter (Supplementary Fig. S1). It is thus possible that this region was responsible for the oligomerisation of PhaC*_Aq_*.

Two small fractions, nevertheless, eluted around 55 and 148 kDa at roughly a ratio of 3:1, which corresponded to monomeric and dimeric PhaC*_Aq_* using a molecular weight of 67.4 kDa (Fig. 1). PhaC*_Cn_* likewise existed predominantly as monomers, and the monomeric form shifted to dimeric in the presence of synthetic (3HB)*_n_*-CoA analogues (*n* = 2–4) with an increase in activity, suggesting that the active form was dimeric [9]. Using the dimeric form of PhaC*_Aq_* for enzyme activity assays, the *K*_m_ and *k*_cat_ values were calculated to be 394 µM and 476 s^−1^, respectively, for the substrate 3HB-CoA. While kinetic constants of different enzymes are hard to compare due to experimental variations, these values were nevertheless comparable to those of PhaC*_Cn_* [19], i.e. 103 µM and 333 s^−1^.

### 3.2. Model of PhaC_Aq_ with a three-branched tunnel

As the C-terminal domain of PhaC*_Aq_* (Glu200–Gln595) shares an even higher identity of 51% with that of PhaC*_Cn_*, a reliable model of PhaC*_Aq_* could be generated from the PhaC*_Cn_* structure for comparison (Fig. 2A). The PhaC*_Aq_* model contained a conserved catalytic triad constituted by Cys319, Asp475 and His505, and was also expected to maintain a similar dimeric interface as many of the residues involved in PhaC*_Cn_* dimerisation were conserved. Some variations, nevertheless, are expected as PhaC*_Aq_* possesses a LID region shorter by five residues (Pro355–Pro414) and lacks the disulphide bond present in PhaC*_Cn_*’s LID region (Cys382–Cys438; Supplementary Fig. S1).

**Fig. 2.**
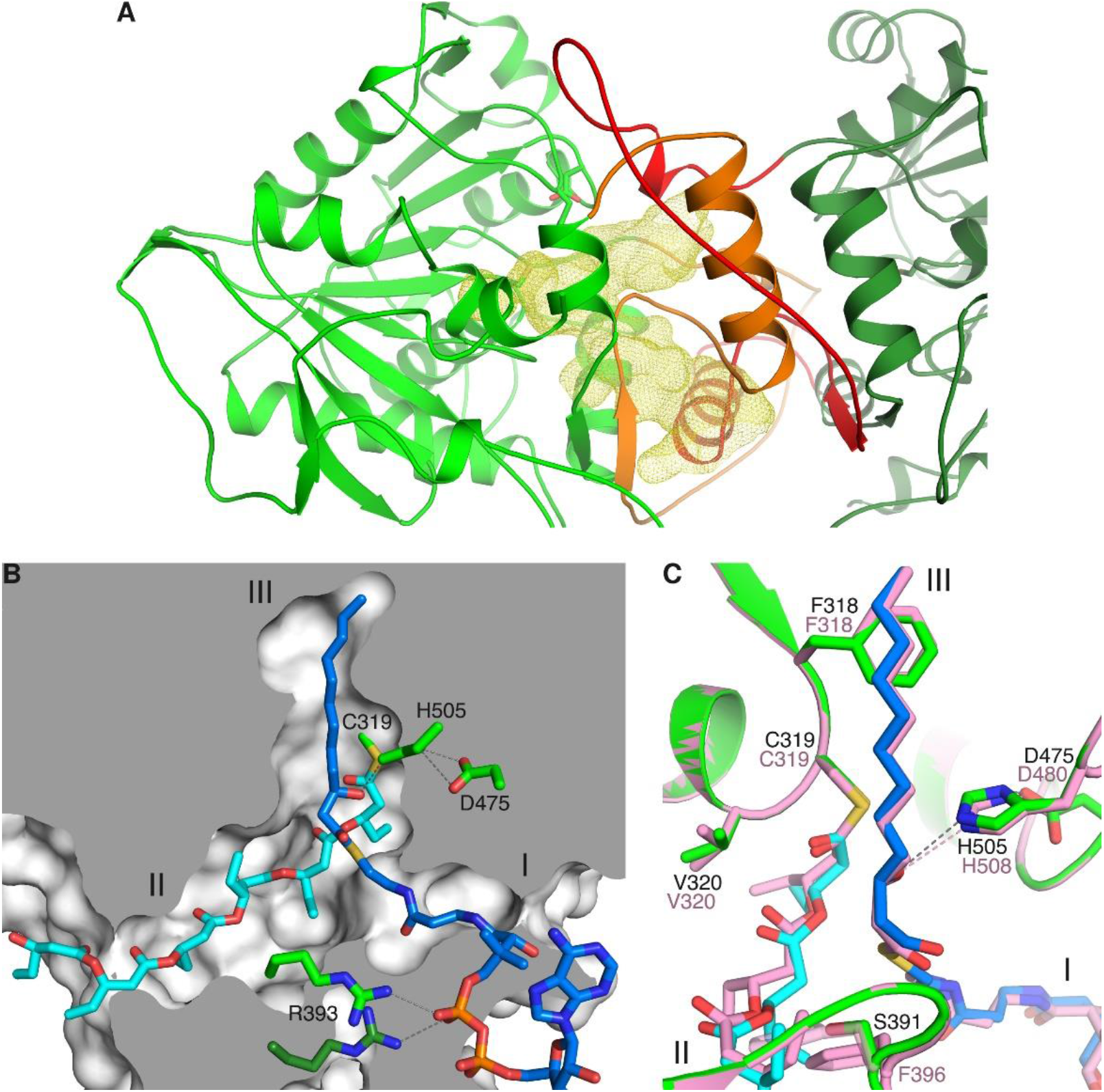
Modelling and docking of class I PhaC*_Aq_*. (**A**) The PhaC*_Aq_* model (green and dark green for the two monomers) contains a large cavity (yellow) at the dimer interface, which is constituted in part by the flexible LID region (orange and red from each monomer). (**B**) The three-branched tunnel is predicted to bind the CoA moiety of a HA-CoA substrate (blue) in branch I and the growing PHA chain (cyan) covalently bonded to Cys319 in branch II, while branch III is capable of accommodating a 3HDD (C12) moiety. (**C**) The catalytic site of PhaC*_Cn_* (pink) also has similar capacity to bind HA-CoA substrates of short to medium chain lengths in branch III, and a PHA chain in branch II after rotating sideways the side chain of its Phe396 (transparent).

The PhaC*_Aq_* model revealed a large cavity around the catalytic triad as observed in PhaC*_Cn_*, which was expected to be the putative substrate binding site (Fig. 2A). Unexpectedly, the cavity further extended to the solvent through an opening, which however was not observed in the PhaC*_Cn_* structure. A tunnel with three branches, I–III, finally emerged after the ligand docking procedures without moving any main chains in the cavity (Fig. 2B). Two of the branches, I and II, traverse the dimer interface, while the catalytic triad is located midway in branch III. The large cavity also observed in PhaC*_Cn_* was formed by both branches I and III, which were proposed respectively for substrate entrance (branch I) and product egress (branch III) [6]. However, the hydrophobic branch III forms a dead end inside the protein core — as noted, it would require large conformational changes around this region to allow a growing PHB chain to pass through [6].

Branch II, on the other hand, is ~19.5 Å long and directly opens up to the solvent, hence serving better as the tunnel for product exit. It is not readily observable in the PhaC*_Cn_* structure mainly because of the bulky side chain of Phe396 at the vertex of the tunnel. Located in the flexible LID region and opposite to the catalytic triad, it replaces the much smaller Ser391 of PhaC*_Aq_* and blocks the entrance to branch II (Fig. 2C). Nevertheless, branch II could be rendered possible in PhaC*_Cn_* by rotating Phe396’s side chain, as well as that of Leu397, sideways. Similarly, a three-branched tunnel was also found in a PhaC*_Cs_* model based on the PhaC*_Cn_* structure, suggesting that branch II is an integral feature of the tunnel probably involved in product passage.

### 3.3. Docking of ligands

The CoA moiety could be docked into branch I in a similar fashion as reported in the PhaC*_Cn_* structure [6], with the strictly conserved Arg393 of PhaC*_Aq_* from both monomers binding to one pyrophosphate group. By rotating mainly the side chains of Trp289, Asp396 and Phe435, meanwhile, a chain of six 3-hydroxyvalerate (3HV; C5) molecules covalently bonded to the catalytic Cys319 was docked into branch II (Fig. 2B). About five 3HV molecules could traverse branch II before exiting through the opening, but the subsites for each monomer were not well defined. In comparison, PhaC*_Cn_*’s branch II is slightly narrower as six of its residues lining the tunnel (Phe352, Ala391, Phe396, Trp436, Tyr437 and Tyr440) are larger than those of PhaC*_Aq_*’s (His352, Gly386, Ser391, Met431, His432 and Phe435).

As both solvent-accessible branches I and II have been delegated the obvious roles in substrate entrance and product exit respectively, the dead-end branch III is left with the possible role in binding the HA moiety of a HA-CoA substrate. Intriguingly, with a length of ~14 Å it seems that PhaC*_Aq_*’s branch III is able to accommodate even HAs of medium chain length. This is also in agreement with the findings that PhaC*_Cn_*, although classified as a class I PhaC that utilises CoA thioesters of short-chain-length HAs, could also incorporate HAs as long as 3-hydroxyoctanoate (3HO; C8) and 3-hydroxydodecanoate (3HDD; C12) [20,21]. Since the residues lining the tunnel of both PhaC*_Aq_* and PhaC*_Cn_* are highly conserved, PhaC*_Aq_* is therefore also expected to be able to bind substrates containing 3HO and 3HDD.

By rotating sideways a few side chains including Leu252 and Met253 without causing any steric strain, a 3HDD moiety indeed could be neatly docked into PhaC*_Aq_*’s branch III whose narrowest point is ~3.6 Å in diameter (Fig. 2B). The 3HDD-CoA substrate could be similarly docked into PhaC*_Cn_* and PhaC*_Cs_*, suggesting that class I PhaCs do share a common structural feature for substrate specificity. However, the lengthy 3HDD monomer would face difficulty in exiting through branch II, which in the present model did not even accommodate short-chain-length HAs comfortably. It would certainly require substantial conformational changes to allow the passage of such medium-chain-length PHA polymers.

Nevertheless, the present PhaC*_Cn_* structure is noted as partially open and could have been induced by the artificial disulphide bond between Cys382 located in the LID region and Cys438, none of which is conserved. This has led to part of its LID region (Leu358–Leu384) adopting an extended loop conformation encircling the opposite subunit, which in the closed-form PhaC*_Cs_* structure actually folded into an α-helix [8]. It is thus probable that dimerisation in a truly functional dimer may differ, with the highly flexible LID region, which makes up a huge area of branches I and II, moving outwards to expand them. A shift of the LID region, for example, may be able to allow the side chain of PhaC*_Aq_*’s Tyr440 to rotate sideways and free up some space for the growing PHA chain.

### 3.4. Modelling of class II PhaC

Due to the comparatively high identity of 43% between their C-terminal domains, a reliable model of a class II PhaC1 from *Pseudomonas aeruginosa*, PhaC1 *_Pa_*, could also be constructed based on the PhaC*_Cn_* structure. As expected, a three-branched tunnel is also observed (Fig. 3A). Compared to class I PhaCs, interestingly, the PhaC1 *_Pa_* model has an elongated branch III of ~22 Å in length. Since the major difference between classes I and II is the ability of class II PhaCs to incorporate HAs of medium and long chain length (> C14), this further lends support to the assumption that the dead-end branch III binds the HA moiety of a HA-CoA substrate.

**Fig. 3.**
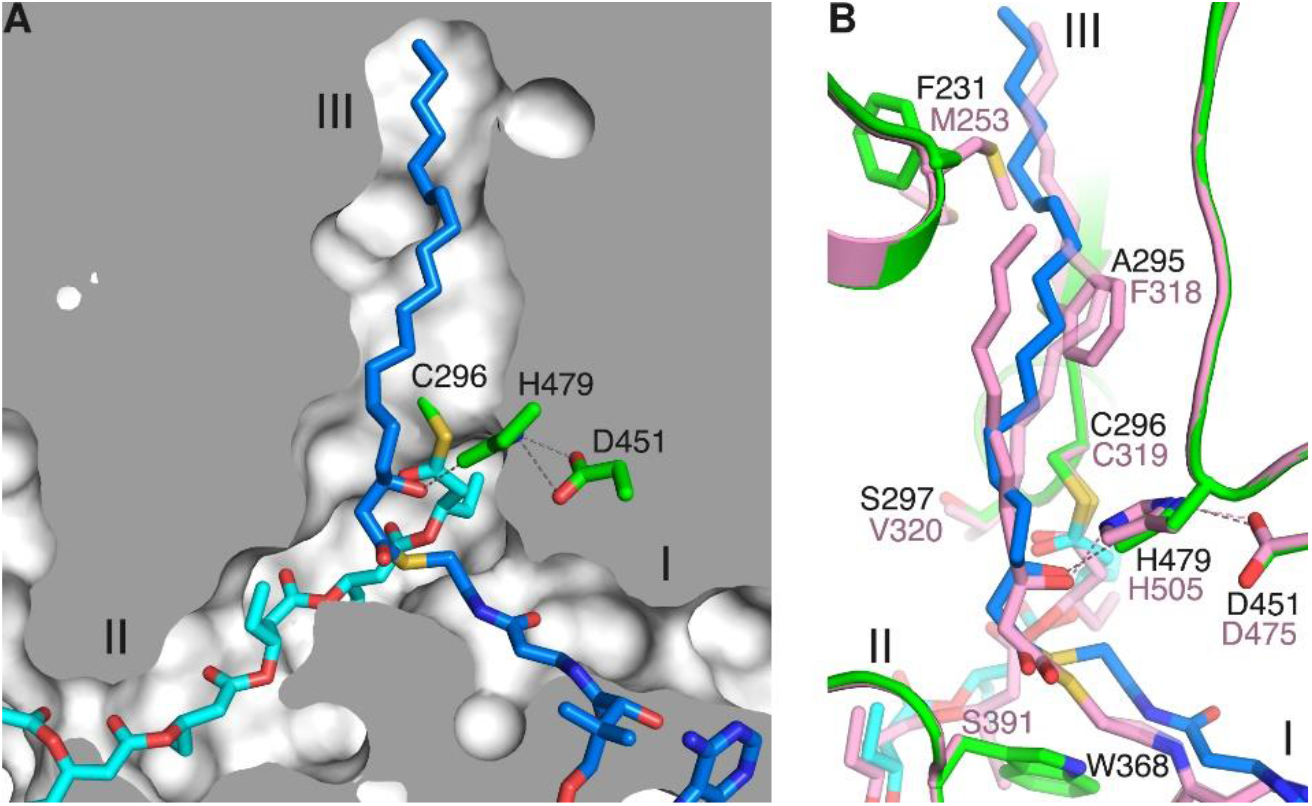
Modelling of class II PhaC1*_Pa_*. (**A**) While branches I and II of the PhaC1*_Pa_* model are similar to those of PhaC*_Aq_*, it has an elongated branch III that can accommodate a C19 moiety (blue). (**B**) Although PhaC*_Aq_* (pink) is unable to bind long-chain-length HAs, changing its Phe318, which is conserved in PhaC1*_Pa_* (green), to Ala in the F318A model (transparent) frees up space for a C18 HA moiety to bind to branch III. The HA moiety moves into the space occupied by Phe318, allowing Met253 to rotate downwards and this in turn further makes way for the HA moiety.

Indeed, a substrate with C19 HA was successfully docked into PhaC1*_Pa_*’s hydrophobic branch III (Fig. 3A). In comparison with class I PhaCs, the highly conserved Phe318 of PhaC*_Aq_* is notably conserved as a much smaller Ala295 in PhaC1*_Pa_*, enabling the latter’s branch III to create the necessary space for a HA of long chain length (Fig. 3B). Likewise, a model of PhaC2 from *P. aeruginosa*, PhaC2*_Pa_*, could also accommodate a C19 HA-CoA substrate. Interestingly, a C18 HA-CoA ligand could be docked into a PhaC*_Aq_* model that had its Phe318 changed to Ala (Fig. 3B), as well as a C17 HA-CoA into a F318A PhaC*_Cn_* model, whose branch III was slightly shorter due to its Met242 replacing PhaC*_Aq_*’s Ile242.

In PhaC1*_Pa_*, the residue located at the tunnel’s vertex is replaced by an even larger Trp368, which however is highly conserved in class II PhaC (Supplementary Fig. S1). Its bulky side chain has rendered branch I narrower for binding the CoA moiety. Branch II of PhaC1*_Pa_*, with a size similar to that of class I PhaCs, is also too small to allow the passage of a growing chain of bulky HAs. This further shows that the present PhaC*_Cn_* structure is not truly representative of a functional dimer. For example, in PhaC1*_Cn_* Val407 from the LID region is sandwiched between two α-helices, but it is replaced by a bulky Trp379 in the PhaC1 *_Pa_* model which in turn dents the neighbouring α-helix.

### 3.5. A common mechanism for PHA elongation

The mechanism for PHA elongation basically involves two steps — transferring a growing (HA)*_n_* chain from the catalytic Cys to a HA-CoA substrate, and then transferring the (HA)_*n*+1_ chain back to the Cys. In other words, the enzyme has to act on two different sites of the substrate — first deprotonation of the HA hydroxyl group, then cleavage of the thioester bond — which will certainly require manoeuvring these two sites into the enzyme’s catalytic site during their respective steps. While the present PhaC*_Cn_* structure represents a partially open form, the catalytic triad as well as branch III are located in the core of the enzyme which undergoes negligible structural changes between the partially open and closed forms, and hence these regions still serve as a solid basis for investigating the catalytic mechanism.

The exact elongation mechanism can be readily inferred by modelling the two respective tetrahedral intermediates into the PhaC*_Aq_* model. The first tetrahedral intermediate is formed after the hydroxyl group of a HA-CoA substrate, deprotonated by His505, nucleophilically attacks the carbonyl group of the Cys319-bonded (HA)*n* chain (Fig. 4A,B). The tetrahedral intermediate is stabilised by the oxyanion hole formed by the backbone NH group of Val320. Protonation by His505 then releases the (HA)_*n*+1_-CoA molecule — in effect transferring the (HA)*_n_* chain from Cys319 to HA-CoA (Fig. 4C).

**Fig. 4.**
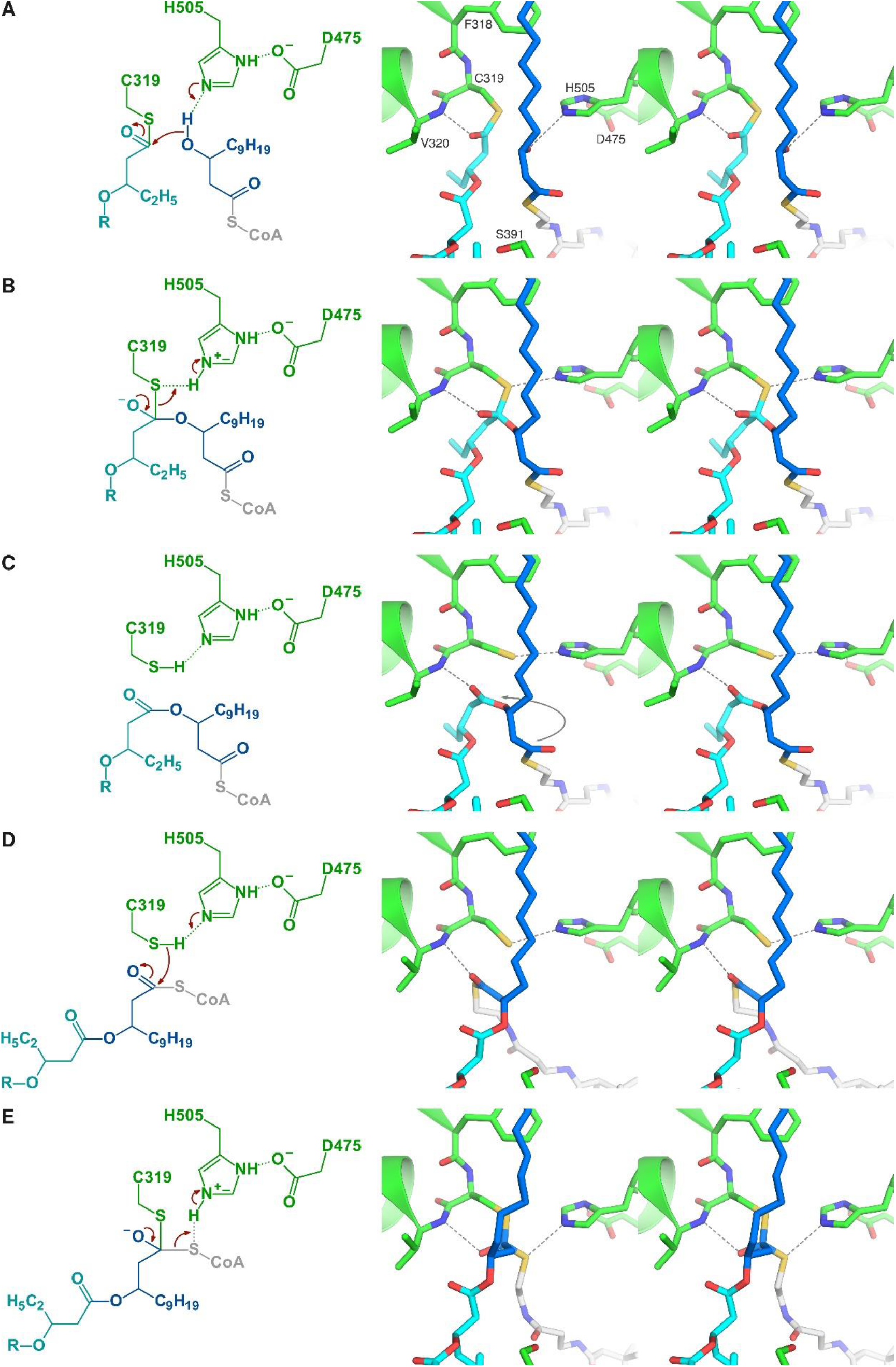
A polymer elongation mechanism for PhaC. The mechanistic scheme (left) and stereo view (right) depict the steps for transferring a HA moiety (blue) from a HA-CoA substrate to Cys319. (**A**) The hydroxyl group of the HA-CoA substrate nucleophilically attacks the carbonyl group of the Cys319-bonded (HA)*_n_* chain (cyan). (**B**) The first tetrahedral intermediate is formed and stabilised by the backbone of Val320. (**C**) Protonation by His505 releases the (HA)_*n*+1_-CoA molecule, whose carbonyl carbon of the scissile thioester bond is then moved into the catalytic site by a ~180° rotation of the HA moiety. (**D**) The same reaction is repeated but with Cys319 acting to carry out the nucleophilic attack on the thioester bond to form the second tetrahedral intermediate. (**E**) CoA is released from the tetrahedral intermediate upon proton donation from His505.

The substrate’s site for the next step, the carbonyl carbon atom of the (HA)_*n*+1_-CoA’s scissile thioester bond, however, is now located more than 6 Å away from Cys319. In order to proceed, the HA moiety of the carbonyl group makes a ~180° rotation to bring the carbonyl carbon into the catalytic site (Fig. 4D). After that basically the same reaction is repeated but with a switch of roles — this time Cys319 carries out the nucleophilic attack on the carbonyl carbon to form the second tetrahedral intermediate, which is again stabilised by Val320 (Fig. 4E). Protonation by His505 finally releases CoA, completing one cycle of chain elongation with the (HA)_*n*+1_ chain transferred back to Cys319.

An empty cavity is observed around Ser391 at the intersection of branches I and II in the PhaC*_Aq_* model. The large side chain of PhaC*_Cn_*’s Phe396, in comparison, is seen better at filling the cavity and hence may act better at segregating and supporting both the HA-CoA substrate and the growing PHA product. PhaC1*_Pa_*’s Trp368, on the other hand, renders branch I narrower for binding the CoA moiety. Upon closer inspection, a Phe residue which precedes PhaC*_Aq_*’s Ser391 by two residues, Phe389, is found highly conserved not only in classes I and II but also in class III PhaCs (Supplementary Fig. S1). The corresponding Phe361 in the closed-form PhaC*_Cs_* structure, interestingly, is located at the intersection. It is possible that this Phe is the actual residue playing the segregating role in a functional dimer, which may rotate sideways upon CoA release to help the extended PHA chain move from branch III to II.

While PhaC is known to exhibit a characteristic lag phase followed by a rapid linear phase with the 3HB-CoA substrate, and the lag phase is known to be reduced by priming PhaC with synthetic (3HB)*_n_*-CoA analogues (*n* = 2–4) [9], the exact basis for both phenomena is unknown. The presence of a three-branched tunnel seems to be able to partially offer a physical explanation. When a 3HB-CoA molecule first enters an intact PhaC molecule, the 3HB moiety can get into either branch III or, by chance, the wider branch II of a functional dimer. The process of tucking the 3HB moiety back into branch III slows down the enzymatic transfer of 3HB to the catalytic Cys. After the enzyme is covalently bonded to one 3HB molecule, any incoming 3HB-CoA molecule may be prevented from entering branch II.

However, the covalently bonded 3HB moiety still has some degree of freedom to vibrate at the intersection and impede the binding of another 3HB-CoA molecule, explaining why the lag phase was still present after PhaC priming. The freedom to vibrate decreases as more 3HB molecules are covalently bonded, in agreement with the observation that priming with a trimer reduced the lag phase more than with a dimer [9]. When the PHB chain has grown to a certain length that allows itself to be tucked nicely into branch II, new incoming 3HB-CoA molecules will then be able to bind directly to the catalytic site and the lag phase disappears. The (3HB)_4_-CoA analogue, on the other hand, was the least efficient primer probably because the tetramer moiety experienced the greatest difficulty in tucking into the empty branch II.

### 3.6. Classification of PhaCs

The class I PhaC structures have provided certain key features for delineating the different classes of PhaCs. A prominent feature is the bulky Phe318 that precedes the catalytic Cys319 of PhaC*_Aq_* (Fig. 2C), which limits the chain length of HA-CoA substrates and is highly conserved as either Phe, Tyr or Trp in class I PhaCs (Fig. 5). This bulky residue is instead conserved strictly as Ala in class II PhaCs, freeing up the necessary space for binding HA-CoA substrates of medium to long chain lengths. While PhaC models for classes III and IV, both of which favour short-chain-length HAs, could not be reliably constructed due to low identity, class IV PhaCs such as those from the *Bacillus* genus likewise possess a conserved Tyr as class I PhaCs, whereas class III PhaCs contain a conserved Ile.

**Fig. 5.**
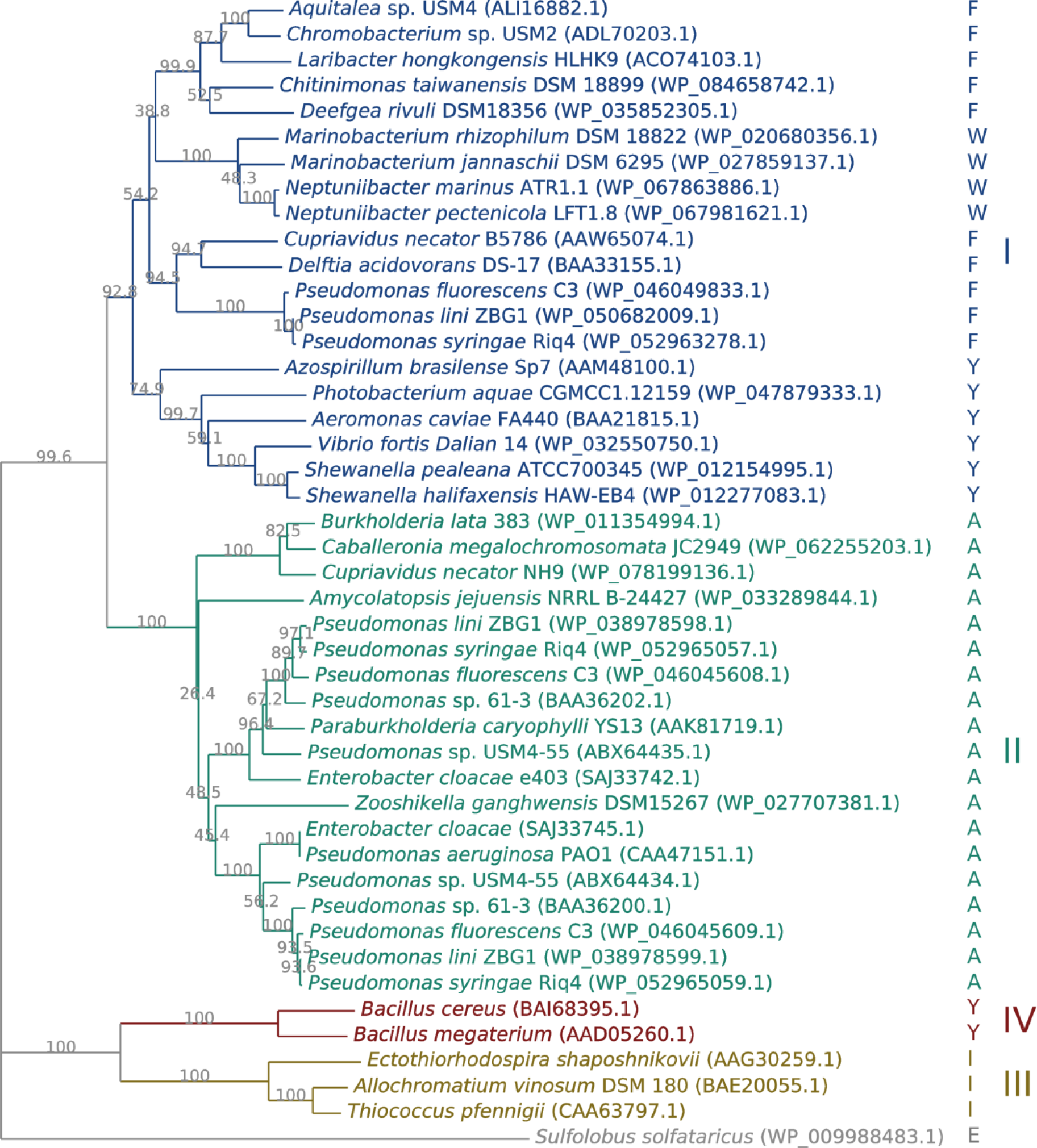
Neighbour-joining phylogenetic tree of PhaCs. PhaC*_Aq_*’s Phe318 is conserved as a bulky Phe, Tyr or Trp in class I (blue), but strictly as the much smaller Ala in class II (green). Bootstrap values shown at the branch points are expressed as percentage of 1,000 replications.

Members from classes III and IV have been reported to also incorporate medium-chain-length HAs like 3HO (C8), for example the class III PhaC from *Thiococcus* (*Thiocapsa*) *pfennigii* [22] and class IV PhaC from *B. cereus* [23]. If classes III and IV PhaCs too share a similar branch III as class I PhaCs, structurally they should be able to bind also HA-CoA substrates of medium chain length. Nevertheless, considering the strictly conserved catalytic triad and a relatively conserved catalytic domain, including PhaC*_Aq_*’s Arg393 and His476 predicted to bind the CoA moiety, among all the classes (Supplementary Fig. S1), classes III and IV PhaCs are expected to share a similar substrate binding mode and a common PHA elongation mechanism.

## 4. Conclusions

While *Aquitalea* PhaC, PhaC*_Aq_*, tends to form large oligomers, a small fraction exists in an equilibrium of monomers and dimers with comparable *K*_m_ and *k*_cat_ values of 394 µM and 476 s^−1^, respectively, for the substrate 3HB-CoA. A PhaC*_Aq_* model, based on the closely related PhaC*_Cn_* crystal structure, has revealed the existence of a three-branched tunnel capable of accommodating HAs as long as 3HDD (C12) in branch III. While branch II for putative product exit is too narrow for HAs such as 3HDD as the model is in a partially open form, the tunnel has nevertheless served as a reasonable basis for explaining the elongation mechanism of PHA, in which a ~180° rotation of the HA moiety is required after the transfer of a (HA)*_n_* chain from the catalytic Cys319 to a HA-CoA. The same mechanism can similarly be applied to the PhaC*_Cn_* structure — confirming the validity of the PhaC*_Aq_* model — as well as to the class II PhaC1*_Pa_* model. In summary, both classes I and II mostly share a common mechanism of enzyme action, and substrate specificity is determined in part by the elongated branch III of class II PhaCs that can accommodate HAs of long chain length.

## Acknowledgments

This work was supported by the APEX Delivering Excellence grant from Universiti Sains Malaysia (1002/PBIOLOGI/910322).

## References

[1] A.J. Anderson, E.A. Dawes, Occurrence, metabolism, metabolic role, and industrial uses of bacterial polyhydroxyalkanoates, Microbiol Rev, 54 (1990) 450-472.

[2] T. Keshavarz, I. Roy, Polyhydroxyalkanoates: bioplastics with a green agenda, Curr Opin Microbiol, 13 (2010) 321-326.

[3] B.H. Rehm, Polyester synthases: natural catalysts for plastics, Biochem J, 376 (2003) 15-33.

[4] P. Schubert, A. Steinbuchel, H.G. Schlegel, Cloning of the *Alcaligenes eutrophus* genes for synthesis of poly-β-hydroxybutyric acid (PHB) and synthesis of PHB in *Escherichia coli*, J Bacteriol, 170 (1988) 5837-5847.

[5] Y. Jia, W. Yuan, J. Wodzinska, C. Park, A.J. Sinskey, J. Stubbe, Mechanistic studies on class I polyhydroxybutyrate (PHB) synthase from *Ralstonia eutropha:* class I and III synthases share a similar catalytic mechanism, Biochemistry, 40 (2001) 1011-1019.

[6] E.C. Wittenborn, M. Jost, Y. Wei, J. Stubbe, C.L. Drennan, Structure of the catalytic domain of the class I polyhydroxybutyrate synthase from *Cupriavidus necator*, J Biol Chem, 291 (2016) 25264-25277.

[7] J. Kim, Y.J. Kim, S.Y. Choi, S.Y. Lee, K.J. Kim, Crystal structure of *Ralstonia eutropha* polyhydroxyalkanoate synthase C-terminal domain and reaction mechanisms, Biotechnol J, 12 (2017).

[8] M.F. Chek, S.Y. Kim, T. Mori, H. Arsad, M.R. Samian, K. Sudesh, T. Hakoshima, Structure of polyhydroxyalkanoate (PHA) synthase PhaC from *Chromobacterium* sp. USM2, producing biodegradable plastics, Sci Rep, 7 (2017) 5312.

[9] J. Wodzinska, K.D. Snell, A. Rhomberg, A.J. Sinskey, K. Biemann, J. Stubbe, Polyhydroxybutyrate synthase: Evidence for covalent catalysis, J Am Chem Soc, 118 (1996) 6319-6320.

[10] L.M. Ng, K. Sudesh, Identification of a new polyhydroxyalkanoate (PHA) producer *Aquitalea* sp. USM4 (JCM 19919) and characterization of its PHA synthase, J Biosci Bioeng, 122 (2016) 550-557.

[11] A. Leaver-Fay, M. Tyka, S.M. Lewis, O.F. Lange, J. Thompson, R. Jacak, K. Kaufman, P.D. Renfrew, C.A. Smith, W. Sheffler, I.W. Davis, S. Cooper, A. Treuille, D.J. Mandell, F. Richter, Y.E. Ban, S.J. Fleishman, J.E. Corn, D.E. Kim, S. Lyskov, M. Berrondo, S. Mentzer, Z. Popovic, J.J. Havranek, J. Karanicolas, R. Das, J. Meiler, T. Kortemme, J.J. Gray, B. Kuhlman, D. Baker, P. Bradley, ROSETTA3: an object-oriented software suite for the simulation and design of macromolecules, Methods Enzymol, 487 (2011) 545-574.

[12] P.D. Adams, P.V. Afonine, G. Bunkoczi, V.B. Chen, I.W. Davis, N. Echols, J.J. Headd, L.W. Hung, G.J. Kapral, R.W. Grosse-Kunstleve, A.J. McCoy, N.W. Moriarty, R. Oeffner, R.J. Read, D.C. Richardson, J.S. Richardson, T.C. Terwilliger, P.H. Zwart, PHENIX: a comprehensive Python-based system for macromolecular structure solution, Acta Crystallogr D Biol Crystallogr, 66 (2010) 213-221.

[13] R.C. Edgar, MUSCLE: multiple sequence alignment with high accuracy and high throughput, Nucleic Acids Res, 32 (2004) 1792–1797.

[14] N. Saitou, M. Nei, The neighbor-joining method: a new method for reconstructing phylogenetic trees, Mol Biol Evol, 4 (1987) 406–425.

[15] S. Kumar, G. Stecher, K. Tamura, MEGA7: Molecular Evolutionary Genetics Analysis Version 7.0 for Bigger Datasets, Mol Biol Evol, 33 (2016) 1870–1874.

[16] T.U. Gerngross, K.D. Snell, O.P. Peoples, A.J. Sinskey, E. Csuhai, S. Masamune, J. Stubbe, Overexpression and purification of the soluble polyhydroxyalkanoate synthase from *Alcaligenes eutrophus*: evidence for a required posttranslational modification for catalytic activity, Biochemistry, 33 (1994) 9311–9320.

[17] Z. Zheng, M. Li, X.J. Xue, H.L. Tian, Z. Li, G.Q. Chen, Mutation on N-terminus of polyhydroxybutyrate synthase of *Ralstonia eutropha* enhanced PHB accumulation, Appl Microbiol Biotechnol, 72 (2006) 896–905.

[18] Y.J. Kim, S.Y. Choi, J. Kim, K.S. Jin, S.Y. Lee, K.J. Kim, Structure and function of the N-terminal domain of *Ralstonia eutropha* polyhydroxyalkanoate synthase, and the proposed structure and mechanisms of the whole enzyme, Biotechnol J, 12 (2017).

[19] S. Zhang, T. Yasuo, R.W. Lenz, S. Goodwin, Kinetic and mechanistic characterization of the polyhydroxybutyrate synthase from *Ralstonia eutropha*, Biomacromolecules, 1 (2000) 244–251.

[20] R.V. Antonio, A. Steinbuchel, B.H. Rehm, Analysis of in vivo substrate specificity of the PHA synthase from *Ralstonia eutropha*: formation of novel copolyesters in recombinant *Escherichia coli*, FEMS Microbiol Lett, 182 (2000) 111–117.

[21] P.R. Green, J. Kemper, L. Schechtman, L. Guo, M. Satkowski, S. Fiedler, A. Steinbuchel, B.H. Rehm, Formation of short chain length/medium chain length polyhydroxyalkanoate copolymers by fatty acid *β*-oxidation inhibited *Ralstonia eutropha*, Biomacromolecules, 3 (2002) 208–213.

[22] M. Liebergesell, F. Mayer, A. Steinbüchel, Analysis of polyhydroxyalkanoic acid-biosynthesis genes of anoxygenic phototrophic bacteria reveals synthesis of a polyester exhibiting an unusal composition, Appl Microbiol Biotechnol, 40 (1993) 292–300.

[23] K.P. Caballero, S.F. Karel, R.A. Register, Biosynthesis and characterization of hydroxybutyrate-hydroxycaproate copolymers, Int J Biol Macromol, 17 (1995) 86–92.

